# GPT-2’s activations predict the degree of semantic comprehension in the human brain

**DOI:** 10.1101/2021.04.20.440622

**Authors:** Charlotte Caucheteux, Alexandre Gramfort, Jean-Rémi King

## Abstract

Language transformers, like GPT-2, have demonstrated remarkable abilities to process text, and now constitute the backbone of deep translation, summarization and dialogue algorithms. However, whether these models encode information that relates to human comprehension remains controversial. Here, we show that the representations of GPT-2 not only map onto the brain responses to spoken stories, but also predict the extent to which subjects understand narratives. To this end, we analyze 101 subjects recorded with functional Magnetic Resonance Imaging while listening to 70 min of short stories. We then fit a linear model to predict brain activity from GPT-2’s activations, and correlate this mapping with subjects’ comprehension scores as assessed for each story. The results show that GPT-2’s brain predictions significantly correlate with semantic comprehension. These effects are bilaterally distributed in the language network and peak with a correlation of R=0.50 in the angular gyrus. Overall, this study paves the way to model narrative comprehension in the brain through the lens of modern language algorithms.

---

**I**n less than two years, language transformers like GPT-2 have revolutionized the field of natural language processing (NLP). These deep learning architectures are typically trained on very large corpora to complete partially-masked texts, and provide a one-fit-all solution to translation, summarization, and question-answering tasks and algorithms (1).

Critically, their hidden representations have been shown to – at least partially – correspond to those of the brain: single-sample fMRI (2–4), MEG (2, 4), and intracranial responses to spoken and written texts (3, 5) can be significantly predicted from a linear combination of the hidden vectors generated by these deep networks. Furthermore, the quality of these predictions directly depends on the models’ ability to complete text (3, 4).

In spite of these achievements, strong doubts subsist on whether language transformers actually encode meaningful constructs (6). When asked to complete “I had $20 and gave $10 away. Now, I thus have $”, GPT-2 predicts “20”^∗^. Similar trivial errors can be observed for geographical locations, temporal ordering, pronoun attribution and causal reasoning. These results have thus led some to argue that such “system has no idea what it is talking about” (7). Thus, how the representations of GPT-2 relate to a human-like understanding remains largely unknown.

Here, we propose to evaluate how the similarity between the brain and GPT-2 vary with semantic comprehension. Specifically, we first compare GPT-2’s activations to the functional Magnetic Resonance Imaging of 101 subjects listening to 70 min of seven short stories, and we quantify this similarity with a “brain score” (ℳ) (8, 9). Second, we evaluate how the brain scores systematically vary with semantic comprehension, as individually assessed by a questionnaire at the end of each story.

## GPT-2’s activations linearly map onto fMRI responses to spoken narratives

To assess whether GPT-2 generates similar representations to those of the brain, we first evaluate, for each voxel, subject and narrative independently, whether the fMRI responses can be predicted from a linear combination of GPT-2’s activations (Figure 1A). We summarize the precision of this mapping with a brain score ℳ: i.e. the correlation between the true fMRI responses and the fMRI responses linearly predicted, with cross-validation, from GPT-2’s responses to the same narratives (cf. Methods). To mitigate fMRI spatial resolution and the necessity to correct each observation by the number of statistical comparisons, we here report either 1) the average brain scores across voxels or 2) the average score within each region of interest (*n* = 314, following an automatic subdivision of Destrieux atlas (10), cf. SI.1). Consistent with previous findings (2, 4, 11, 12), these brain scores are significant over a distributed and bilateral cortical network, and peak in middle- and superior-temporal gyri and sulci, as well as in the supra-marginal and the infero-frontal cortex (2, 4, 11) (Figure 1B).

**Fig. 1.**
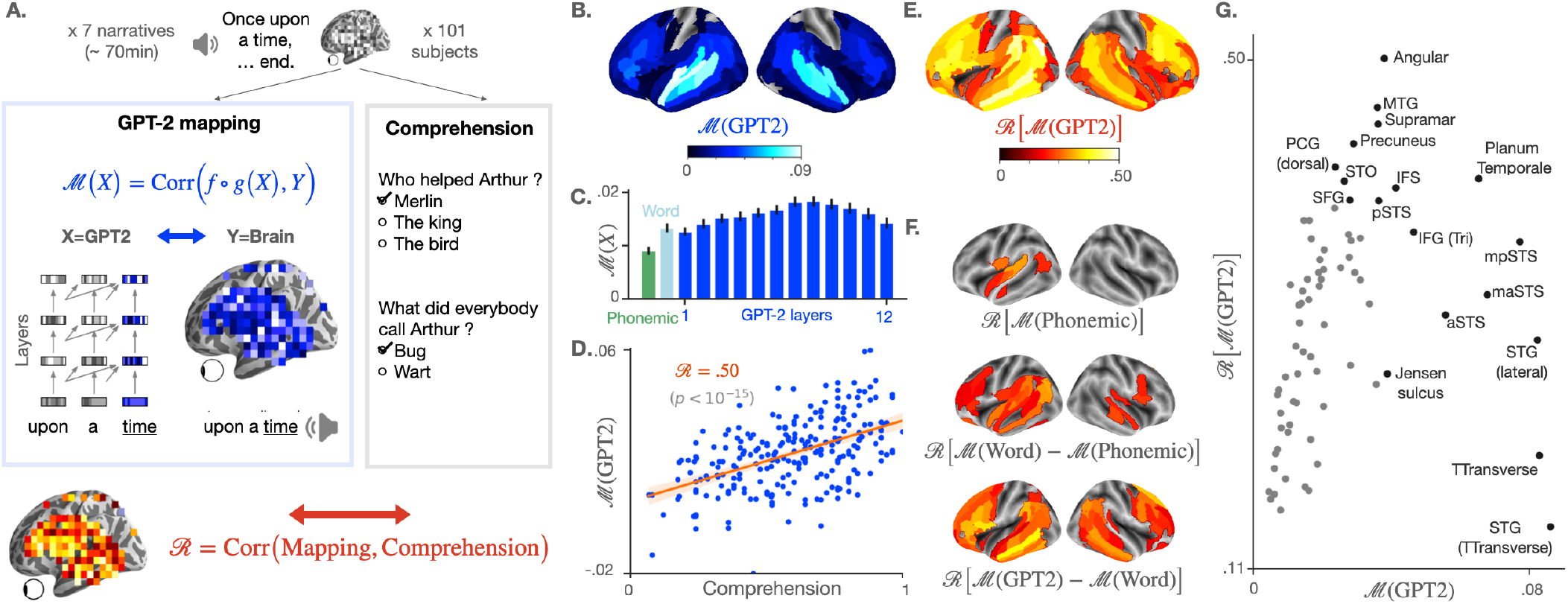
**A**. 101 subjects listen to narratives (70 min of unique audio stimulus in total) while their brain signal is recorded using functional MRI. At the end of each story, a questionnaire is submitted to each subject to assess their understanding, and the answers are summarized into a comprehension score specific to each (narrative, subject) pair (grey box). In parallel (blue box on the left), we measure the mapping between the subject’s brain activations and the activations of GPT-2, a deep network trained to predict a word given its past context, both elicited by the same narrative. To this end, a linear spatio-temporal model (*f* ∘ *g*) is fitted to predict the brain activity of one voxel *Y*, given GPT-2 activations *X* as input. The degree of mapping, called “brain score” is defined for each voxel as the Pearson correlation between predicted and actual brain activity on held-out data (blue equation, cf. Methods). Finally, we test the correlation between the comprehension scores of the subjects and their corresponding brain scores using Pearson’s correlation (red equation). A positive correlation means that the representations shared across the brain and GPT-2 are key for the subjects to understand a narrative. **B**. Brain scores (fMRI predictability) of the activations of the eighth layer of GPT-2. Scores are averaged across subjects, narratives, and voxels within brain regions (142 regions in each hemisphere, following a subdivision of Destrieux Atlas (10), cf. SI.1). Only significant regions are displayed, as assessed with a two-sided Wilcoxon test across (subject, narrative) pairs, testing whether the brain score is significantly different from zero (threshold: .05). **C**. Brain scores, averaged across fMRI voxels, for different activation spaces: phonological features (word rate, phoneme rate, phonemes, tone and stress, in green), the non-contextualized word embedding of GPT-2 (“Word”, light blue) and the activations of the contextualized layers of GPT-2 (from layer one to layer twelve, in blue). The error bars refer to the standard error of the mean across (subject, narrative) pairs (n=237). **D**. Comprehension and GPT-2 brain scores, averaged across voxels, for each (subject, narrative) pair. In red, Pearson’s correlation between the two (denoted ℛ), the corresponding regression line and the 95% confidence interval of the regression coefficient. **E**. Correlations (ℛ) between comprehension and brain scores over regions of interest. Brain scores are first averaged across voxels within brain regions (similar to B.), then correlated to the subjects’ comprehension scores. Only significant correlations are displayed (threshold: .05). **F**. Correlation scores (ℛ) between comprehension and the subjects’ brain mapping with phonological features (M(Phonemic) (i), the share of the word-embedding mapping that is not accounted by phonological features ℳ(Word) − ℳ(Phonemic) (ii) and the share of the GPT-2 eighth layer’s mapping not accounted by the word-embedding ℳ(GPT2) − ℳ(Word) (iii). **G**. Relationship between the average GPT-2-to-brain mapping (eighth layer) per region of interest (similar to B.), and the corresponding correlation with comprehension (ℛ, similar to D.). Only regions of the left hemisphere, significant in both B. and E. are displayed. In black, the top ten regions in terms of brain and correlation scores (cf. SI.1 for the acronyms). Significance in D, E and F is assessed with Pearson’s p-value provided by SciPy^†^. In B, E and F, p-values are corrected for multiple comparison using a False Discovery Rate (Benjamin/Hochberg) over the 2 *×* 142 regions of interest.

By extracting GPT-2 activations from multiple layers (from layer one to layer twelve), we confirm that middle layers best map onto the brain (Figure 1C), as seen in previous studies (2, 4, 11). For clarity, the following analyses focus on the activations extracted from the *eighth* layer, i.e. GPT-2’s most “brain-like” layer (Figure 1B).

## GPT-2’s brain predictions correlate with semantic comprehension

Does the linear mapping between GPT-2 and the brain reflect a fortunate correspondence (4)? Or, on the contrary, does it reflect similar representations of high-level semantics? To address this issue, we correlate these brain scores to the level of comprehension of the subjects, assessed for each subject-story pair. On average across all voxels, this correlation reaches ℛ = 0.50 (*p* < 10^−15^, Figure 1D, as assessed across subject-story pairs with the Pearson’s test provided by SciPy). This correlation is significant across a wide variety of the bilateral temporal, parietal and prefrontal cortices typically linked to language processing (Figure 1E). Together, these results suggest that the shared representations between GPT-2 and the brain reliably vary with semantic comprehension.

## Low-level processing only partially accounts for the correlation between comprehension and GPT-2’s mapping

Low-level speech representations typically vary with attention (13, 14), and could thus, in turn, influence down-stream comprehension processes. Consequently, one can legitimately wonder whether the correlation between comprehension and GPT-2’s brain mapping is simply driven by variations in low-level auditory processing. To address this issue, we evaluate the predictability of fMRI given low-level phonological features: the word rate, phoneme rate, phonemes, stress and tone of the narrative (cf. Methods). The corresponding brain scores correlate with the subjects’ understanding (ℛ = 0.17, *p* < 10^−2^) but less so than the brain scores of GPT-2 (Δℛ = 0.32). These low-level correlations with comprehension peak in the left superior temporal cortex (Figure 1F). Overall, this result suggests that the link between comprehension and GPT-2’s brain mapping may be partially explained by – but not reduced to – the variations of low-level auditory processing.

## The reliability of high-level representations best predict comprehension

Is the correlation between comprehension and GPT-2’s mapping driven by a *lexical* process and/or by an ability to meaningfully *combine* words? To tackle this issue, we compare the correlations obtained from GPT-2’s word embedding (i.e. layer 0) to those obtained from GPT-2’s eighth layer, i.e. a contextual embedding. On average across voxels, the correlation with comprehension is 0.12 lower with GPT-2’s word embedding than with its contextual embedding. An analogous analysis, comparing word embedding to phonological features is displayed in 1F. Strictly lexical effects (word-embedding *versus* phonological) peak in the superior-temporal lobe and in pars triangularis. By contrast, higher-level effects (GPT-2 eighth layer *versus* word-embedding) peak in the superior-frontal, posterior superior-temporal gyrus, in the precuneus and in both the triangular and opercular parts of the inferior frontal gyrus – a network typically associated with high-level language comprehension (4, 15–19).

## Comprehension effects are mainly driven by individuals’ variability

The variability in comprehension scores could result from exogeneous factors (e.g. some stories may be harder to comprehend than others for GPT-2) and/or from endogeneous factors (e.g. some subjects may better understand specific texts because of their prior knowledge). To address this issue, we fit a linear mixed model to predict comprehension scores given brain scores, specifying the narrative as a random effect (cf. SI.1). The fixed effect of brain score (shared across narratives) is highly significant: *β* = 0.04, *p* < 10^−29^, cf. SI.1). However, the random effect (slope specific to each single narrative) is not (*β* < 10^−2^, *p* > 0.11). We also replicate the main analysis (Figure 1D) within each single narrative: the correlation with comprehension reaches 0.76 for the ‘sherlock’ story and is above 0.40 for every story (cf. SI.1). Overall, these analyses confirm that the link between GPT-2 and semantic comprehension is mainly driven by subjects’ individual differences in their ability to make sense of the narratives.

## Discussion

Our analyses reveal a positive correlation between semantic comprehension and the degree to which GPT-2 maps onto brain responses to spoken narratives.

These results strengthen and complete prior work on the brain bases of semantic comprehension. In particular, previous studies have used inter-subject brain correlation to reveal the brain regions associated with understanding (17). For example, Lerner et al. recorded subjects’ fMRI while they listened to normal texts or texts scrambled at the word, sentence or paragraph level, in order to parametrically manipulate their level of comprehension (15). The corresponding fMRI signals correlated across subjects in the primary and secondary auditory areas even when the input was scrambled below the lexical level. By contrast, fMRI signals also became correlated in the bilateral infero-frontal and temporo-parietal cortex when the scrambling was either not performed, or performed at the level of sentences and paragraphs. Our results are consistent with this hierarchical organization, and thus make an important step towards the development of a cerebral model of narrative comprehension.

The relationship between GPT-2’s representations and human comprehension remains to be qualified. First, although highly significant, our brain scores are relatively low (2, 9, 17). This phenomenon likely results from a mixture of different elements: i) we ran our analyses across *all* voxels to avoid selection biases, which automatically reduces the average effect sizes and ii) we report the results without correcting for a noise ceiling (cf. SI.1), as our pilot analyses suggest that such noise-ceiling can greatly vary depending on how it is implemented (i.e. fit from mean across subjects, from all or on voxels etc). Second, the correlation between semantic comprehension and GPT-2’s mapping is robust (*p* < 10^−15^) but far from perfect (***R*** = 0.50). Such correlation thus indicates that the modeling of brain responses with GPT-2 does not *fully* account for the variation in comprehension. While this result is expected (7), our study provides a promising framework to evaluate the extent to which deep language models represent and understand texts like we do.

Finally, our results suggest that the neural bases of comprehension relate to the *high-level* representations of deep language models. While the mapping of phonological features and word embeddings do correlate with comprehension, GPT-2’s contextual embeddings provides brain maps that more reliably predict comprehension (Figure 1F). The superiority of contextual-embedding in predicting comprehension suggests that i) GPT-2 encodes features supporting comprehension and ii) our finding are not solely driven by low- or mid-level processing (13, 14). These elements remain solely based on correlations, however. The factors that *causally* influence comprehension, ranging from prior knowledge, attention and language complexity should be explicitly manipulated in future work.

Overall, the present study strengthens and clarifies the similarity between the brain and deep language models, repeatedly observed in the past three years (2–4, 11, 20). Together, these findings reinforce the relevance of deep language models in unraveling the neural bases of narrative comprehension.

## Materials and Methods

Our analyses rely on the “Narratives” dataset (21), composed of the brain signals, recorded using fMRI, of 345 subjects listening to 27 narratives.

### Narratives and comprehension score

Among the 27 stories of the dataset, we selected the seven stories for which subjects were asked to answer a comprehension questionnaire at the end, and for which the answers varied across subjects (more than ten different comprehension scores across subjects), resulting in 70 minutes of audio stimuli in total, from four to 19 minutes per story (Figure 2). Questionnaires were either multiple-choice, fill-in-the blank, or open questions (answered with free text) rated by humans (21). Here, we used the comprehension score computed in the original dataset which was either a proportion of correct answers or the sum of the human ratings, scaled between 0 and 1 (21). It summarizes the comprehension of one subject for one narrative (specific to each (narrative, subject) pair).

**Fig. 2.**
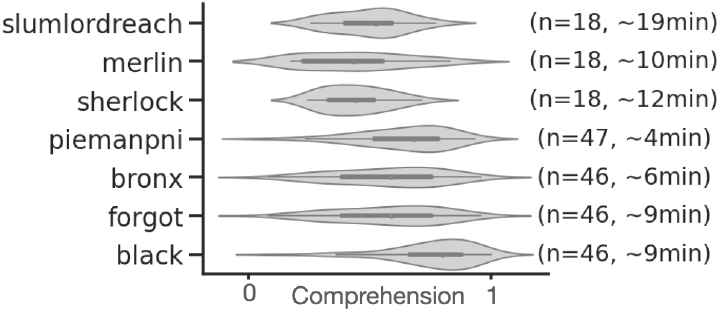
For each of the seven narratives: number of subjects (*n*), distribution of comprehension scores across subjects and length of the narrative.

### Brain activations

The brain activations of the 101 subject who listened to the selected narratives were recorded using fMRI, as described in (21). As suggested in the original paper, pairs of (subject, narrative) were excluded because of noisy recordings, resulting in 237 pairs in total.

### GPT-2 activations

GPT-2 (1) is a high-performing neural language model trained to predict a word given its previous context (it does not have access to succeeding words), given millions of examples (e.g Wikipedia texts). It consists of multiple Transformer modules (twelve, each of them called “layer”) stacked on a non-contextual word embedding (a look-up table that outputs a single vector per vocabulary word) (1). Each layer ℓ can be seen as a nonlinear system that takes a sequence of *w* words as input, and outputs a contextual vector of dimension (*w, d*), called the “activations” of layer ℓ (*d* = 768). Intermediate layers were shown to better encode syntactic and semantic information than input and output layers (22), and to better map onto brain activity (2, 4). Here, we show that the *eighth* layer of GPT-2 best predicts brain activity 1C. We thus select the eighth layer of GPT-2 for our analyses. Our conclusions remain unchanged with other intermediate-to-deep layers of GPT-2 (from 6^*th*^ to 12^*th*^ layers).

In practice, the narratives’ transcripts were formatted (replacing special punctuation marks such as “–” and duplicated marks “?.” by dots), tokenized using GPT-2 tokenizer and input to the GPT-2 pretrained model provided by Huggingface ^‡^. The representation of each token is computed separately using a context window a 1024. For instance, to compute the representation of the third token of the story, we input GPT-2 with the third, second and first token, and then extract the activations corresponding to the third token. To compute the representation of a token *w*_*k*_ at the end of the story, GPT-2 is input with this token combined with the 1,023 preceding tokens. Then, we extract the activations corresponding to *w*_*k*_. The procedure results in a vector of activations of size (*w, d*) with *w* the number of tokens in the story and *d* the dimensionality of the model. There are fewer fMRI scans than words. Thus, the activation vectors between successive fMRI measurements are summed to obtain one vector of size *d* per measurement. To match the fMRI measurements and the GPT-2 vectors over time, we used the speech-to-text correspondences provided in the fMRI dataset (21).

### Linear mapping between GPT-2 and the brain

For each (subject, narrative) pair, we measure the mapping between i) the fMRI activations elicited by the narrative and ii) the activations of GPT-2 (layer nine) elicited by the same narrative. To this end, a linear spatiotemporal model is fitted on a train set to predict the fMRI scans given the GPT-2 activations as input. Then, the mapping is evaluated by computing the Pearson correlation between predicted and actual fMRI scans on a held out set *I*:

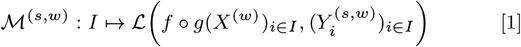

With *f* ∘ *g* the fitted estimator (g: temporal and f: spatial mappings), ℒ Pearson’s correlation, *X*^(*w*)^ the activations of GPT-2 and *Y* ^(*s,w*)^ the fMRI scans of subjects *s*, both elicited by the narrative *w*.

In practice, *f* is a ℓ_2_-penalized linear regression. We follow scikit-learn implementation^§^ with ten possible regularization parameters log-spaced between 10^−1^ and 10^8^, one optimal parameter per voxel and leave-one-out cross-validation. *g* is a finite impulse response (FIR) model with 5 delays, where each delay sums the activations of GPT-2 input with the words presented between two TRs. For each (subject, narrative) pair, we split the corresponding fMRI time series into five contiguous chunks using scikit-learn cross-validation. The procedure is repeated across the five train (80% of the fMRI scans) and disjoint test folds (20% of the fMRI scans). Pearson correlations are averaged across folds to obtain a single score per (subject, narrative) pair. This score, denoted ℳ(*X*) in Figure 1A, measures the mapping between the activations space *X* and the brain of one subject, elicited by one narrative.

### Phonological features

To account for low-level speech processing, we computed the alignment (Eq. (1)) between the fMRI brain recordings *Y* and phonological features *X*: the word rate (of dimension *d* = 1, the number of words per fMRI scan), the phoneme rate (*d* = 1, the number of phonemes per fMRI scan) and the concatenation of phonemes, stresses and tones of the words in the stimuli (categorical feature, *d* = 117). The latter features are provided in the original Narratives database (21), and computed using Gentle^¶^ forced-alignment algorithm.

### Significance

Significance was either assessed by using either (i) a second-level Wilcoxon test (two-sided) across subject-narrative pairs, testing whether the mapping (one value per pair) was significantly different from zero (Figure 1B), or (ii) by using the first-level Pearson p-value provided by SciPy^‖^ (Figure 1D-G). In Figure 1B, E, F, p-values were corrected for multiple comparison (2 × 142 ROIs) using False Discovery Rate (Benjamin/Hochberg)^∗∗^.

## Supporting Information (SI)

### Brain parcellation

In Figure 1B, E, and F, we used a subdivision of the parcellation from Destrieux Atlas (10). Regions with more than 400 vertices were split into smaller regions (so that each regions contains less than 400 vertices). The original parcellation consists of 75 regions per hemisphere. Our custom parcellation consists in 142 regions per hemisphere.

In Figure 1G, we use the original parcellation for simplicity, and the following acronyms:

| Acronym | Definition |
| --- | --- |
| STG / STS | Superior temporal gyrus / sulcus |
| aSTS | Anterior STS |
| maSTS | Mid-anterior STS |
| mpSTS | Mid-posterior STS |
| pSTS | Posterior STS |
| Angular / Supramar | Angular / Supramarginal inferior parietal gyrus |
| MTG / MTS | Medial temporal gyrus / sulcus |
| SFG / SFS | Superior frontal gyrus / sulcus |
| IFG / IFS | Inferior frontal gyrus / sulcus |
| Tri / Op | Pars triangularis / opercularis (IFG) |
| TTransverse | Temporal transverse sulcus |
| PCG | Posterior cingulate gyrus |
| STO | Temporo-occipital lateral sulcus |

### Mixed-effect model

Not all subjects listened to the same stories. To check that the ℛ scores (correlation between comprehension and brain mapping) were not driven by the narratives and questionnaires’ variability, a linear mixed-effect model was fit to predict the comprehension of a subject given its brain mapping scores, specifying the narrative as a random effect. More precisely, if *w*_*i*_ ∈ ℝ corresponds to the mapping scores of the *i*^*th*^ subject that listened to the story *w*, and 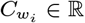 refers to the comprehension scores, we estimate the fixed effect parameters 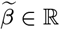 and 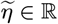 (shared across narratives), and the random effect parameter *β*_*w*_ ∈ ℝ and *η*_*w*_ ∈ ℝ (specific to the narrative *w*) such that:

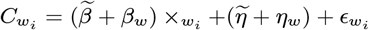

with 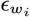 a vector of i.i.d normal errors with mean 0 and variance *σ*^2^. In practice, we use the statsmodels^††^ implementation of linear mixed-effect models. Significance of the coefficients were assessed with a t-test, as implemented in statsmodels.

### Replication across single narratives

To further support that the ℛ were not driven by the narratives’ variability, we replicate the analysis of Figure 1D within single narratives. In Figure 3, we show that correlation scores between brain scores and comprehension scores are positive for each of the seven narratives.

**Fig. 3.**
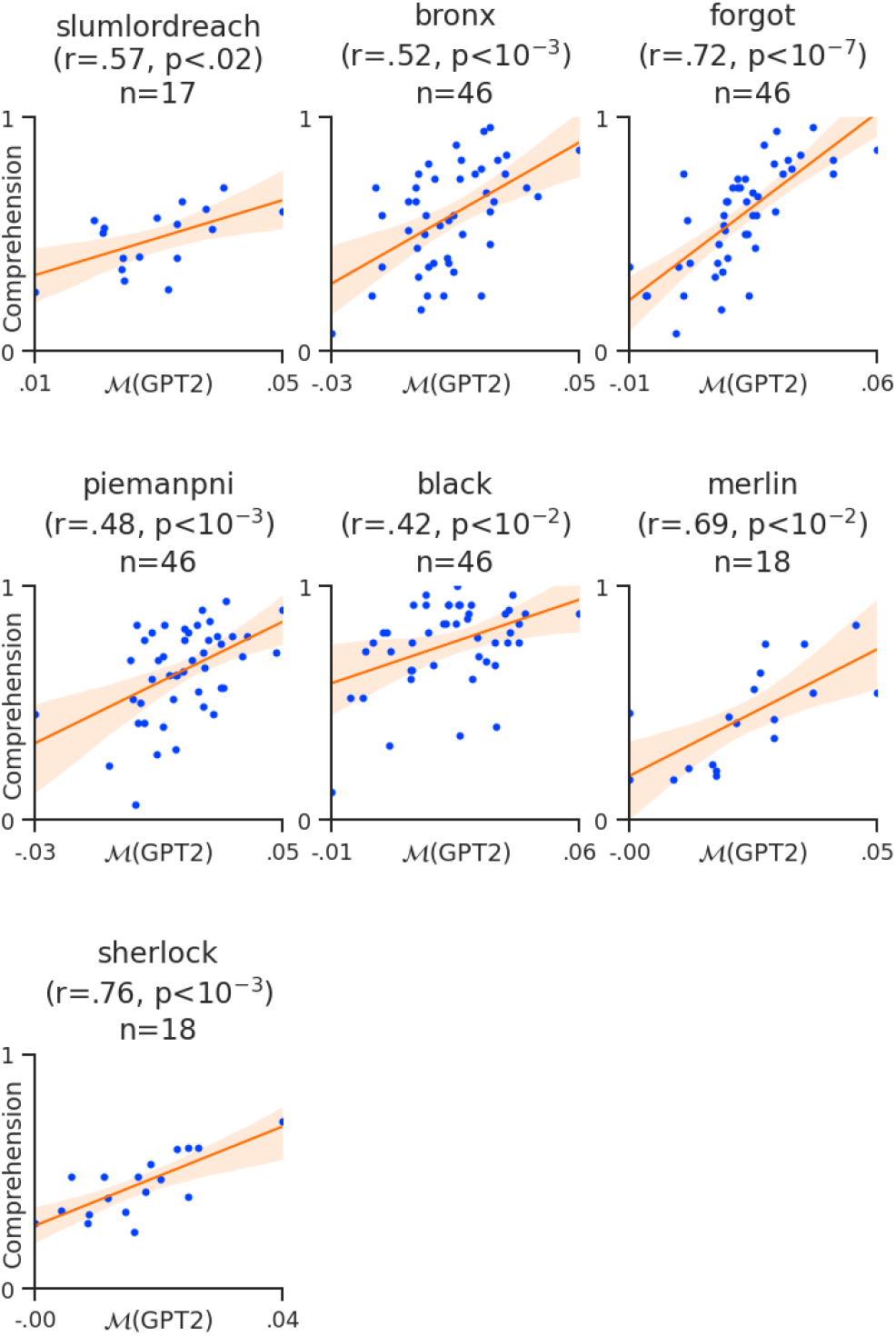
Replication within single narratives. Same as Figure 1D for each single narrative.

### Noise Ceiling Estimates

fMRI recordings are inherently noisy. Thus, we estimate an upper bound of the best brain score that can be obtained given the level of noise in the Narrative dataset. To this end, for each (subject, narrative) pair, we linearly map the fMRI recordings, not with the GPT-2 activations, but with the average fMRI recordings of the other subjects who listened to that narrative. More precisely, we use the exact same setting as in Eq. (1), but we predict *Y* ^(*s*)^, not from *g*(*X*) (GPT-2’s features after temporal alignment, of size *n*_times_ × *n*_dim_), but from the mean of the other subject’s brains 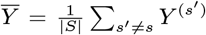 (of size *n*_times_ × *n*_voxels_). This score is called the noise ceiling for the (subject, narrative) pair. The noise ceilings for each brain region are displayed in Figure 4, and correspond to upper bounds of the brain scores displayed in Figure 1B.

**Fig. 4.**
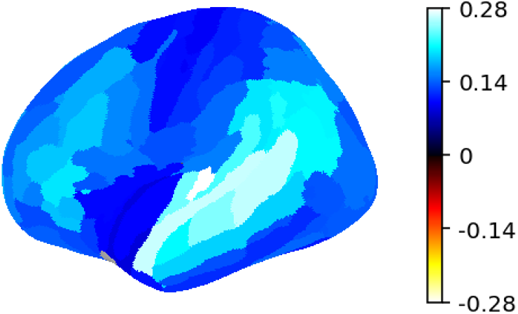
Noise ceiling estimates. Noise ceilings averaged across subjects, narratives and voxels within each region of interest. They are upper bounds of the brain scores in Figure 1B.

## Footnotes

‡ https://github.com/huggingface/transformers

§ https://scikit-learn.org/

¶ https://github.com/lowerquality/gentle

‖ https://www.scipy.org/

∗∗ https://mne.tools/

†† https://www.statsmodels.org/

